# Inflammatory and oxidative status in European captive black rhinoceroses: a link with Iron Overload Disorder?

**DOI:** 10.1101/2020.03.26.009811

**Authors:** Hanae Pouillevet, Nicolas Soetart, Delphine Boucher, Rudy Wedlarski, Laetitia Jaillardon

## Abstract

Iron Overload Disorder (IOD) is a syndrome developed by captive browsing rhinoceroses like black rhinoceroses (*Diceros bicornis*) in which hemosiderosis settles in vital organs while free iron accumulates in the body, potentially predisposing to various secondary diseases. Captive grazing species like white rhinoceroses (*Ceratotherium simum*) do not seem to be affected. The pro-oxidant and pro-inflammatory properties of iron, associated with the poor antioxidant capacities of black rhinoceroses, could enhance high levels of inflammation and oxidative stress leading to rapid ageing and promoting diseases. In this prospective study, 15 black (BR) and 29 white rhinoceroses (WR) originating from 22 European zoos were blood-sampled and compared for their iron status (serum iron), liver/muscle biochemical parameters (AST, GGT, cholesterol), inflammatory status (total proteins, protein electrophoresis) and oxidative stress markers (SOD, GPX, dROMs). Results showed higher serum iron and liver enzyme levels in black rhinoceroses (P<0.01), as well as higher GPX (P<0.05) and dROM (P<0.01) levels. The albumin/globulin ratio was lower in black rhinoceroses (P<0.05) due to higher α_2_-globulin levels (P<0.001). The present study suggests a higher inflammatory and oxidative profile in captive BR than in WR, possibly in relation to iron status. This could be either a consequence or a cause of iron accumulation, potentially explaining rapid ageing and various diseases. Further investigations are needed to assess the prognostic value of the inflammatory and oxidative markers in captive black rhinoceroses, particularly for evaluating the impact of reduced-iron and antioxidant-supplemented diets.

## Introduction

Black rhinoceroses *(Diceros bicornis,* BR) are browser rhinoceroses found in eastern and southern Africa. The three extant wild subspecies, *i.e.* south-western BR (*D. b.* ssp. *bicornis*), eastern BR (*D. b.* ssp. *michaeli*) and southern-central BR (*D. b.* ssp. *minor*), are considered vulnerable to critically endangered by the International Union for Conservation of Nature (IUCN) [1]. Recently, international collaboration enabled the translocation of five BR from three European zoos to Akagera National Park in Rwanda, to diversify the gene-pool and enable healthy population growth in the park [2].

Still, *ex situ* conservation of BR in zoological institutions remains challenging because captive individuals develop several diseases not described in wild BR [3], including hemolytic anemia, hepatopathy, ulcerative dermatopathy and Iron Overload Disorder (IOD). The latter is a syndrome that is being exponentially described in captive BR [4–7], but is not reported in wild BR [4,8–10] nor in grazer rhinoceroses such as white rhinoceroses (*Ceratotherium simum*, WR), whether they be captive or wild. This syndrome is a form of iron storage disease due to free iron accumulation within the organism, leading to hemosiderosis and subsequent hemochromatosis in vital organs, potentially enhancing organ failure in BR [10,11]. The longer the time spent in captivity, the more severe the disease [12]. Currently, the main hypothesis to explain captive BR’s susceptibility to iron accumulation is a discrepancy between the captive and the natural diet, which may lead to increased availability of iron in the captive diet [6,11,13–15].

In humans, hemochromatosis is considered as an inflammatory disease [16] with increased oxidative stress [17]. Oxidative stress has severe consequences on health through high tissue and cellular toxicity [16–21] thus participating in cancer formation [18,22] and promoting secondary diseases and rapid ageing. Even finely regulated in the healthy state, non-transferrin bound iron (NTBI, also called free iron) is able to accept and donate electrons readily thus enhancing the formation of free radicals and consequently oxidative stress [17,23]. Under pathological conditions, iron and superoxide metabolisms are strongly interactive and can exacerbate the toxicity of the other, leading to a self-sustained and ever-increasing spiral of cytotoxic and mutagenic events [17]. This interaction has already been suggested in a BR since they seem to experience a high susceptibility to oxidative stress compared to other mammals [24–29] due to their impaired antioxidant capacities that appeared to be compounded by iron overload.

In the present study, the authors hypothesized that inflammation and oxidative stress may be implicated in the pathogenesis of IOD in captive BR, making this syndrome a potential common denominator to various diseases described in captivity in this species. This study was thus designed to compare inflammation status and oxidative stress levels in relation to iron status in captive BR and WR, the latter being a species theoretically unaffected by IOD.

## Materials and methods

### Study population

Fifteen BR from 8 European zoos (nine females and six males, aged 5-33 yr-old) and 29 WR from 14 European zoos (18 females and six males, five unknown, aged 4-46 yr-old) were prospectively included between May 2017 and May 2018. Within the 4/15 BR for which the medical background was available, one was reported with infertility, one with ulcerative dermatitis, and one with joint pain or arthritis, whereas the last one had not experienced any infection or illness to the veterinarian’s knowledge. Regarding the 21/29 WR for which the medical background was available, 3/21 were reported with arthritis/joint pain, 1/21 with carpal tumour, 1/21 with allergic conjunctivitis and rhinitis, 4/21 with suspected infertility, and the 13/21 remaining had not experienced any infection or illness to the veterinarians’ knowledge. None of these captive BR and WR was reported to receive iron chelators.

### Sample collection, processing and analysis

This prospective study was validated and supported upstream by coordinators of both BR and WR European Association of Zoos and Aquaria’s European Endangered Species Programmes. Each rhinoceros was blood sampled opportunistically by zoo veterinarians, whether during an anaesthesia procedure or a medical training session that was planned to occur independently from the present study. Three millilitre blood samples were collected whether from the auricular or radial vein, both in a heparin and a dry tube. The dry tube was settled for two hours, then centrifuged (1.733 g for five minutes) and the serum was transferred into a clean dry tube. Within 7 days after sampling, both heparinized whole blood and serum tubes were sent to the veterinary laboratory (LDHVet-LabOniris, Nantes, France). Median delay between the sampling and the reception by the lab was 3 days [range from 1 to 7 days] including 2 days of shipment [range from 1 to 3 days]. Directly after reception, whole blood and serum were aliquoted and stored at −20°C until analyses were performed.

All the serum biochemistry was performed using an automated biochemistry analyser (RX Daytona, Randox Laboratories, Crumlin, County Antrim, United Kingdom), unless indicated otherwise. Iron status was evaluated through serum iron measurement (ferrozine colorimetric method). As the liver and muscle are reported as the first tissues suffering from IOD in BR [30], the hepatic and muscular functions were investigated through the measurement of the following parameters: aspartate aminotransferase (AST, L-aspartate/α-oxoglutarate as substrate), gamma glutamyltransferase (GGT, L-γ-glutamyl-3-carboxy-4-nitroanilide/glycylglycine as substrate), cholesterol (cholesterol esterase/oxidase method) and creatine kinase (CK, creatine phosphate as substrate). Inflammation status was evaluated by measuring total serum protein (TP, biuret method) and agarose gel electrophoretic albumin and globulin fractions (albumin and globulins including α_1_-, α_2_-, β- and γ-globulin fractions, agarose gel method (Hyrys2, Hydragel; Sebia, Evry, France). Globulin fractions were determined according to recent published data from Hooijberg and colleagues [31]. α_1_-globulins refers to α_1_-a and α_1_-b fractions, and β-globulins refers to β1 and β2 fractions. The albumin:globulin ratio (A/G) was calculated. Finally, oxidative stress was assessed through the measurement of superoxide dismutase (SOD) and glutathione peroxidase (GPX) activities (colorimetric methods with RANSOD and RANSEL test kits respectively), and reactive oxygen metabolites (dROMs, Diacron Reactive Oxygen Metabolites d-ROMs test, Diacron Laboratories Grosseto, Italy).

No heparinized whole blood was received for one of the BR in which GPX and SOD could thus not be measured. A blood clot was observed in the heparin tube of one WR. As a consequence, GPX, SOD and dROMs could not be measured. CK levels were measured after all the other analyses: as such, this information is missing for two BR and three WR for which serum was lacking. Finally, in one of the WR, protein electrophoresis showed aberrant results which remained unexplained and it was thus excluded from the results.

### Statistical analyses

All statistical analyses were carried out using R-Studio software version 1.1.442 [32]. The data were pooled across species, and basic descriptive statistics, including arithmetic mean, median, standard deviation (SD), minimum and maximum, were obtained for each parameter. As no variable was normally distributed, non-parametric statistical tests were used. Mann Whitney test was performed for all the quantitative parameters (age, iron, AST and CK activities, cholesterol, total proteins, albumin, SOD and GPx) in order to compare BR and WR. Graphic assessment of the SOD and GPx data showed a potential discriminant threshold (>1500 g/Hb and >300 U/gHb, respectively) between the 2 populations of rhinos. As a result, these data were transformed into categorical variables (“Normal” versus “High”) for the statistical analysis. For GGT and dROMs values, as some results were out of the linearity range of the analyser (10 rhinos had GGT values <8 U/L and 13 had dROMs > 1,000 UCARR), a 2 level categorization was also performed using a graphically determined threshold (13 U/L for GGT and 800 UCARR for dROMs). Chi-squared tests were used to compare BR and WR for sex, CatGGT, CatdROMs, CatSOD and CatGPX. Statistical significance was set at P < 0.05.

## Results

Measured parameters for both European captive BR and WR are listed in Table 1. Sex distribution did not differ significantly between BR and WR (P=0.32) as well as age (median 21 years [5–33] versus 16 years [4–46], respectively) (P = 0.32). Serum iron was significantly higher (P < 0.01) in BR (median 42.0 [26.6-58.9] μmol/L) compared to WR (28.0 [11.1-58.4] μmol/L) (Fig 1). Regarding liver and muscular function, AST activity (96 [72-152] versus 71 [12-178] U/L for BR and WR, respectively) (Fig 1), CatGGT (100% “High” *i.e* GGT>13 U/L versus 20.8% for BR and WR, respectively) and CK (323 [199-945] versus 196 [129-697] U/L for BR and WR, respectively) were significantly higher (P< 0.01) in BR compared to WR. Total proteins were significantly higher (P = 0.03) in BR (84 [65-92] g/L) compared to WR (78 [70-95] g/L) (Fig 1). A/G ratio was significantly lower (P = 0.01) in BR (0.56 [0.31-0.83]) compared WR (0.73 [0.20-1.07]) because of significantly higher (P < 0.001) levels of α_2_-globulin (16.210.2-21.9] g/L and 11.2 [6.7-15.8] g/L for BR and WR, respectively) (Figs 1 and 2) Finally,regarding oxidative stress assessment, CatdROMs was significantly higher (P < 0.001) in BR (93.3 % “High” *i.e* dROMs > 800 UCARR) in comparison to WR (32.1% “High” *i.e* dROMs > 800 UCARR), as well as CatGPX (P = 0.047; 64.3% versus 32.1% of “High” *i.e.* GPX > 300 U/gHb, for BR and WR, respectively).

**Table 1.** Results for parameters measured in European captive black and white rhinoceroses for the evaluation of iron status, hepatic and muscular function, inflammation status and oxidative stress levels.^a^.

|  | Black rhinoceros |  |  |  |  |  |  | White rhinoceros |  |  |  |  |  |  |
| --- | --- | --- | --- | --- | --- | --- | --- | --- | --- | --- | --- | --- | --- | --- |
|  | Mean | SD | Min | Median | Max | “High” | n | Mean | SD | Min | Median | Max | “High” | n |
| Age (years) | 19 | 9.2 | 5 | 21 | 33 |  | 15 | 16 | 10.0 | 4 | 16 | 46 |  | 26 |
| Serum iron (μmol/L) | 42.2 | 10.7 | 26.6 | 42.0<sup>b</sup> | 58.9 |  | 15 | 29.8 | 9.9 | 11.1 | 28.0 | 58.4 |  | 29 |
| AST (U/L) | 104 | 21.4 | 72 | 96<sup>b</sup> | 152 |  | 15 | 77 | 33.4 | 12 | 71 | 178 |  | 29 |
| CK (U/L) | 37<br>9 | 201.7 | 199 | 323<sup>b</sup> | 945 |  | 13 | 246 | 142.9 | 129 | 196 | 697 |  | 26 |
| GGT (U/L) | 25 | 8.1 | 14 | 24 | 45 |  | 15 | c | c | < 8 | 10 | 20 |  | 29 |
| CatGGT (%) |  |  |  |  |  | 100<sup>b</sup> | 15 |  |  |  |  |  | 20.8 | 29 |
| Cholesterol (g/L) | 0.7 | 0.2 | 0.5 | 0.7 | 1.3 |  | 15 | 0.8 | 0.4 | 0.4 | 0.7 | 2.6 |  | 29 |
| TP (g/L) | 82.0 | 8.3 | 65.0 | 84.0<sup>b</sup> | 92.0 |  | 15 | 78.2 | 5.7 | 70.0 | 78.0 | 95.0 |  | 29 |
| A/G ratio | 0.57 | 0.16 | 0.31 | 0.56<sup>b</sup> | 0.83 |  | 15 | 0.71 | 0.17 | 0.20 | 0.73 | 1.07 |  | 28 |
| Albumin (g/L) | 29.4 | 6.6 | 15.3 | 31.5 | 38.0 |  | 15 | 32.1 | 5.9 | 13.3 | 32.4 | 40.4 |  | 28 |
| α<sub>1</sub>-globulin (g/L) | 6.2 | 1.2 | 4.0 | 6.3 | 7.8 |  | 15 | 6.1 | 0.8 | 4.6 | 6.1 | 7.6 |  | 28 |
| α<sub>2</sub>-globulin (g/L) | 15.8 | 3.0 | 10.2 | 16.2<sup>b</sup> | 21.9 |  | 15 | 11.3 | 1.9 | 6.7 | 11.2 | 15.8 |  | 28 |
| β-globulin (g/L) | 16.4 | 3.4 | 11.9 | 17.1 | 21.8 |  | 15 | 16.5 | 2.8 | 13.4 | 15.9 | 26.1 |  | 28 |
| γ-globulin (g/L) | 14.0 | 4.2 | 8.1 | 12.9 | 23.7 |  | 15 | 12.4 | 2.9 | 7.7 | 12.4 | 22.5 |  | 28 |
| SOD (U/g Hb) | 1737 | 340.3 | 1130 | 1730 | 2400 |  | 14 | 1636 | 481.3 | 900 | 1505 | 3300 |  | 28 |
| CatSOD (%) |  |  |  |  |  | 50 |  |  |  |  |  |  | 71.4 |  |
| GPX (U/g Hb) | 324 | 134.2 | 104 | 346 | 560 |  | 14 | 240 | 113.8 | 47 | 239 | 423 |  | 28 |
| CatGPX (%) |  |  |  |  |  | 64.3<sup>b</sup> |  |  |  |  |  |  | 32.1 |  |
| dROMs (U CARR) | c | c | 416 | 978 | > 1,000 |  | 15 | c | c | 380 | 686 | > 1,000 |  | 28 |
| CatdROMs (%) |  |  |  |  |  | 93.3<sup>b</sup> | 15 |  |  |  |  |  | 32.1 | 28 |

<sup>a</sup> A/G ratio indicates albumin/globulin ratio; CatGGT, GGT values categorized as “Normal” or “High” (> 13 U/L); CatSOD, SOD values categorized as “Normal” or “High” (> 1,500 U/gHb); CatGPX, GPX values categorized as “Normal” or “High” (> 300 U/g/Hb); CatdROMs, dROMs categorized as “Normal” or “High” (> 800 UCARR).
<sup>b</sup> Statistically significant (P < 0.05) differences between black rhinoceroses and white rhinoceroses.
c Not calculated because of values out of measurable assay range.

**Fig. 1.**
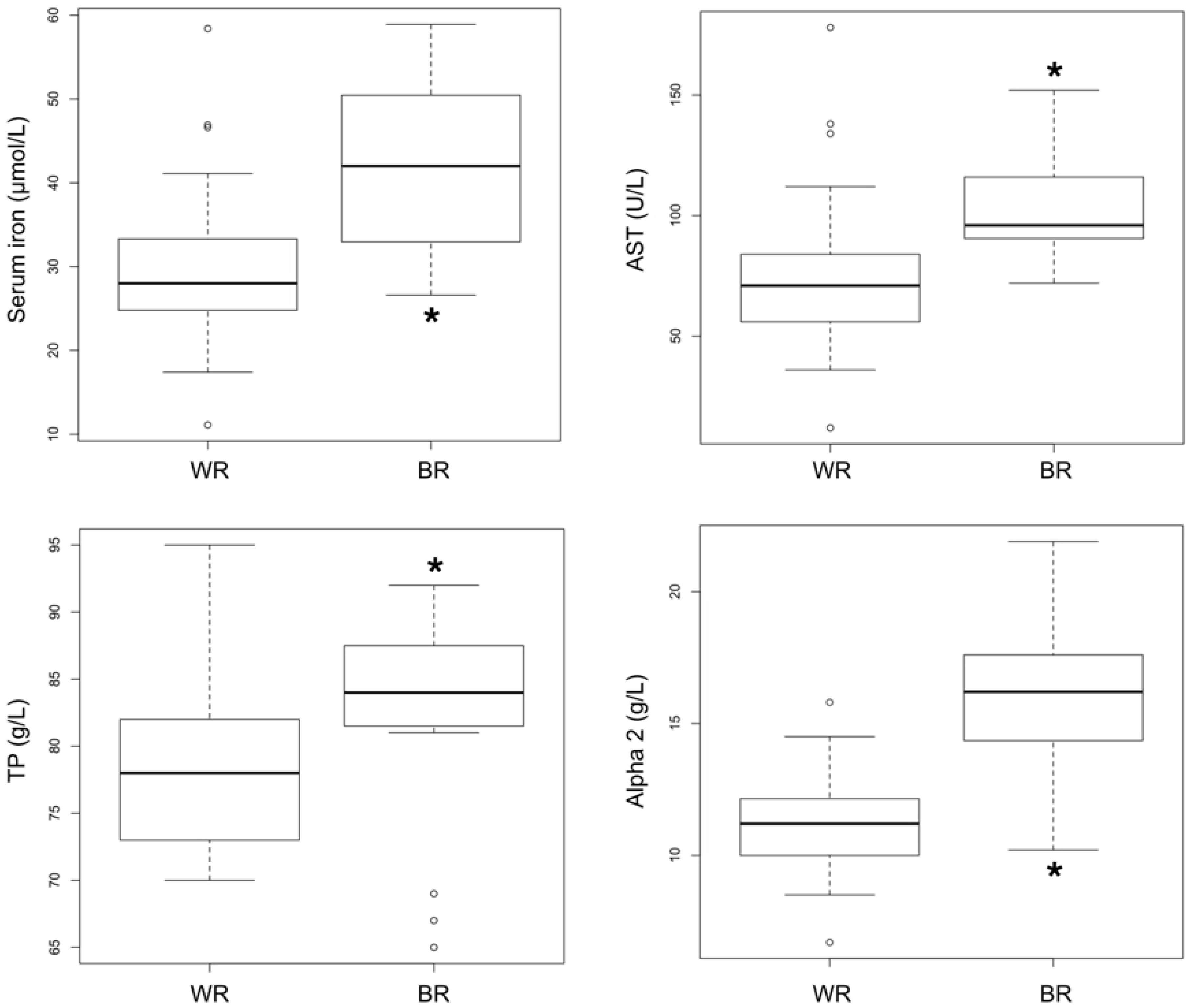
Boxplots showing the quartiles and outliers of serum iron, AST, TP and α_2_-globulins results in 15 European captive black rhinoceroses (BR) and 29 white rhinoceroses (WR). Values were significantly different between the species in all cases (P < 0.05).

**Fig. 2.**
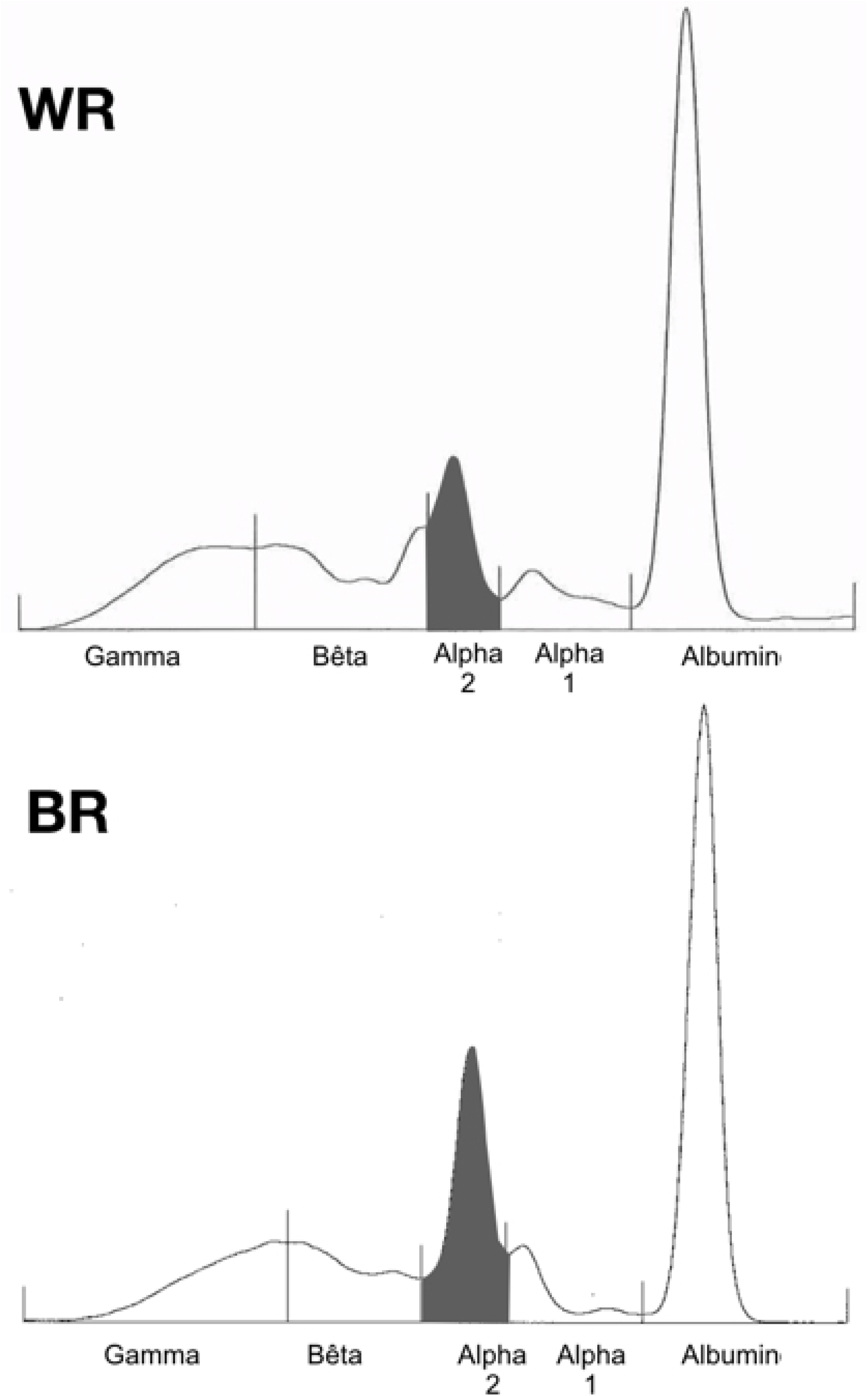
Serum protein electrophoreses showing an example of the serum protein distribution in a black rhinoceros (BR, total proteins 92 g/L, α_2_ globulins 27 g/L) and a white rhinoceros (WR, total proteins 75 g/L, α_2_ globulins 9.6 g/L). Note the increased size of the α_2_ globulin region in the BR compared to the WR (grey areas).

## Discussion

This study showed that European captive black rhinoceroses exhibited higher serum iron concentration and higher inflammatory and oxidative status than captive white rhinoceroses. Taken together, these findings suggest that BR could be predisposed to iron accumulation probably leading to IOD and enhancing inflammatory and oxidative states.

Serum iron values found for the European captive BR included in the present study were very similar to those available in the literature regarding European [10] and American [14,33] captive BR. Serum iron levels being higher in captive BR compared to captive WR has already been reported [8,33]. As such, these results could confirm a predisposition of captive European BR to develop IOD, as previously described [34]. Even if liver biopsy remains the gold standard for definitive diagnosis of iron overload syndromes [35] and has already been performed in a live captive BR confirming diffuse hemosiderosis [36], this procedure is technically challenging due to the animal’s size, the depth of the liver, the difficulty of ultrasound and the skin thickness. Despite having been used in several studies [8,37], ferritin is not specific for iron overload syndromes [38] and is reported as a poor biomarker for IOD progression in Sumatran rhinoceroses, another species of browser rhinoceros [39]. As a consequence, serum iron, TIBC (Total Iron Binding Capacity) measurement and subsequent calculation of Transferrin Saturation may be currently the best tools for guiding *ante-mortem* diagnosis and prognosis of IOD in captive BR as suggested by several studies [10,33,34,36] and should be included in regular blood tests when checking for the health status of captive BR. Measurement of the Total Iron Binding Capacity (TIBC) was intended in the present study through a direct method (TIBC_2, colorimetric method, RX Daytona, Randox Laboratories, Crumlin, County Antrim, United Kingdom), but results were aberrant with many values of TIBC greater than the serum iron concentration, leading to a calculated Transferrin Saturation greater than 100%, which is physiologically not possible. Briefly, in this method, iron is first removed from transferrin through acidification of the sample and a known amount of iron is added. All iron is then complexed and coloured with an iron-binding dye, and absorbance is measured. A second reagent is then added, making the pH rise, resulting in a large increased affinity of transferrin for iron. The observed decrease in absorbance of the coloured dye-iron complex is directly proportional to the TIBC of the serum [40]. We hypothesized that transferrin from rhinoceroses was not as pH-sensitive as human transferrin, and that pH variations were not adequate to accurately measure TIBC in this species, leading to underestimated values. Other methods, such as evaluation of unbound iron binding capacity (UIBC) might be more suitable to calculate Transferrin Saturation in rhinoceroses, as reported in humans [41,42].

Liver function was assessed through AST and GGT measurements, as recommended for domestic horses [43], especially when investigating hemochromatosis [44,45]: indeed, the horse is considered as a very good domestic model animal for rhinoceroses [46,47]. In the present study, increased values of AST in BR compared to WR could have two main causes, including compromised liver function as suggested by increased GGT, and muscular lesions as suggested by increased CK. Molenaar and colleagues reported similar results for GGT in European captive BR [10]. IOD could be implicated in both hepatic and muscular dysfunctions. Indeed, the liver is the first site of hemochromatosis when IOD develops in BR, which impairs its function [16,30]. Iron deposition in muscles has been reported in BR affected by IOD [30]. As hemochromatosis, the latest stage of hemosiderosis, is a progressive and irreversible process with fibrosis that eventually leads to hepatic cirrhosis or carcinoma and fatality in humans [48–50], hepatic biochemistry parameter measurements could be useful prognostic factors in BR affected by IOD.

Higher TP and decreased A/G ratio due to an increase of the α_2_-globulin fraction were observed in captive BR compared to WR, highly suggestive of an increased inflammatory state [51], as it is described in humans affected by hemochromatosis [16,52]. In this study, selecting captive WR instead of free-ranging BR as the negative control precisely aimed at limiting the captivity bias since captivity may be pro-inflammatory by itself [37,53]. Serum protein electrophoretic results of the captive WR included in this study showed some variations in comparison to healthy free-ranging WR [31], further underlining the interest of selecting captive WR as a control group in order to limit the captivity bias. Obesity is described for being pro-oxidative and pro-inflammatory [54], but no study to date investigates the consequences of overweight in rhinoceroses. It would have been relevant to determine whether the European captive BR population did not exhibit significant obesity compared to the European captive WR at the time of the study. However, even if a body condition scoring system has been proposed for BR and Indian rhinoceroses [55,56], no such system exists for WR. The main proteins migrating in the α_2_ region on serum protein electrophoresis include haptoglobin, α_2_-macroglobulin, ceruloplasmin and serum amyloid A (SAA). Since Smith and colleagues have already reported that haptoglobin levels were not significantly different between 10 BR and 20 WR kept in captivity [8], it may not play an important role in the BR’s α_2_-globulin increase in the present study. α_2_-macroglobulin inhibits numerous endogenous proteases and acts as a transport protein for cytokines and growth factors [57]. Increased α_2_-macroglobulin is favored by inflammatory states, such as diabetes mellitus in humans [57]. Ceruloplasmin levels increase with inflammation in humans [58]. Among many roles, this ferroxidase helps to reduce circulating free ferrous iron [17]. As a consequence, it could be hypothesized that ceruloplasmin levels may increase in BR with iron accumulation in response to inflammation and high levels of circulating free iron. As such, its measurement could be of interest in future studies on IOD in BR. Finally, SAA, that increases during inflammation, was reported to be significantly higher in captive BR compared to wild BR [53]. As a result, an increase in α_2_-macroglobulin, ceruloplasmin and SAA, which all suggest inflammation, could explain the increase in α_2_-globulin observed in the captive BR in this study. This finding suggests that IOD could be a chronic inflammatory disease like hemochromatosis is in humans [16]. Inflammation is known for inducing tissue iron storage through hepcidin stimulation and as a consequence for progressively leading to hemochromatosis, thus aggravating iron overload syndromes [59]. Inflammation could thus be a cause and a consequence of IOD in BR, making IOD a potential self-sustaining disease if no iron overload or inflammatory state management is undertaken, with regular phlebotomies as previously described [60].

dROMs are hydroperoxydes, meaning reactive oxygen metabolites that allow to directly evaluate oxidative stress levels [61,62]. SOD and GPX are antioxidant enzymes whose activity measurement indirectly allows to quantifying the response to oxidative stress [63,64]. In the present study, significantly higher levels of GPX and dROMs were observed in captive BR compared to WR, suggesting a higher degree of oxidative stress in captive BR. Increase in oxidative stress levels in captive BR could be due to their lower antioxidant capacities [25,28,29], captivity itself [37,53] and/or a self-sustaining process in which oxidative stress disrupts antioxidant defences [63]. High levels of oxidative stress can predispose to diseases and rapid ageing [17] and should be taken into consideration for husbandry, health care and more globally *ex-situ* conservation of endangered species like the BR. Oxidative stress is reported to worsen iron overload syndromes like iron overload cardiomyopathies in humans [65], which themselves favour reactive oxygen metabolite formation and thus oxidative stress increase [17]. IOD could then be self-sustained through the oxidative stress induced, leading to a vicious circle. However, no significant difference was found concerning SOD. One explanation may be that SOD is not as implicated in BR’s antioxidant defences as in other mammals, making this biomarker possibly not adapted to this species. Regardless, due to low animal numbers, these results should be interpreted with caution.

The main limit of the present study is the postulate that all captive BR may be affected by IOD whereas captive WR are not, without performing a liver biopsy for a definitive diagnosis of IOD. As such, the hypothesized link between increased inflammation and oxidative stress levels and the development of IOD in captive BR need further evaluations to be confirmed. Pre-analytical homogeneity is sub-optimal in the present study. Indeed, food, husbandry and medical management including treatments administered to captive BR and WR could not be controlled. Even if none of their diet included iron chelators, the amount of iron in each diet was not assessed. Also, each zoo institution that collected the blood samples performed the first steps of the processing *i.e.* centrifugation and serum transfer to a new dry tube. The detailed collection and processing protocol submitted to the institutions aimed at limiting this bias. After reception, all samples were aliquoted, frozen and stored at the laboratory for varying durations before they were analysed: this was done in order to group the tests. In humans, most common biochemical analytes show adequate stability in serum following 30 days of storage at −20°C [66], which was the temperature used in the present study for aliquot storage. Nevertheless, it cannot be ruled out that those pre-analytical variations may have compromised the reliability of some blood test results, as no study is available on the stability of such parameters in rhinoceros blood samples.

## Conclusions

Findings of the present study suggest that captive Black Rhinoceroses exhibit higher iron concentrations, higher inflammatory status and higher oxidative stress levels than captive White Rhinoceroses. Taken together, these findings suggest that BR could be predisposed to iron accumulation probably leading to IOD and enhancing inflammatory and oxidative states. Both oxidative stress and inflammation may favour secondary diseases, rapid ageing, and aggravation of IOD, tissue hemochromatosis and organ fibrosis. Thus, efforts to control IOD progression in this endangered species when kept in captivity should be continued, whether through iron level control in the diet or through regular therapeutic phlebotomies. Further investigations are needed to assess the prognostic value of the inflammatory and oxidative markers in captive BR, particularly for evaluating the impact of reduced-iron and antioxidant-supplemented diets.

## Acknowledgments

The authors would like to thank the 22 European zoos that have participated in this prospective study by sending blood samples of their hosted black and white rhinoceroses, and LDHVet - LabOniris for performing the blood analyses.

